# Severe traumatic brain injury temporally affects cerebral blood flow, endothelial cell phenotype, and cilia

**DOI:** 10.1101/2024.11.19.623875

**Authors:** Ankan Gupta, Zachary Bice, Vaya Chen, Yiliang Chen, Anthony J. Veltri, Chien-Wei Lin, Xialong Ma, Amy Y. Pan, Rahima Zennadi, Sean P. Palecek, Ashraf M. Mohieldin, Surya M. Nauli, Ramani Ramchandran, Kevin R. Rarick

## Abstract

**Background:** Previous clinical work suggested that altered cerebral blood flow (CBF) in severe traumatic brain injury (sTBI) correlates with poor executive function and clinical outcome. However, the molecular consequences of altered CBF on endothelial cells (ECs) and their blood flow-sensor organelle called cilia are not known.

**Methods:** We performed laser speckle contrast imaging, single cell isolation, and single cell RNA sequencing (scRNAseq) after sTBI in a closed skull, linear impact mouse model. Validation of select ciliary target protein changes was performed using flow cytometry. Additionally, *in vitro* experiments modeled the post-injury hypoxic environment to evaluate the effect on cilia protein ARL13B in human brain microvascular ECs.

**Results:** We detected immediate reductions in CBF that were sustained for at least 100 minutes in both impacted and non-impacted sides of the brain. Our scRNAseq data detected heterogeneity in the brain cortex-derived EC cluster and demonstrated that two of five unique EC sub-clusters changed their relative proportions post-sTBI. Consistent with flow changes, we identified multiple genes associated with the fluid shear stress pathway that were significantly differentially expressed in brain ECs post-injury. Also, ECs displayed activation of ischemic pathway as early as day 1 post-injury, and enrichment of hypoxia pathway at day 7 and 28 post-injury. *Arl13b* ciliary gene expression was lost on day 1 in ECs cluster and remained lost for the entire course of the injury. We validated the loss of cilia protein ARL13B specifically from brain ECs as early as day 1 post-injury and detected the protein in the peripheral blood of the injured mice. We also determined that hypoxia could induce loss of ARL13B protein from cultured ECs.

**Conclusions:** In severe TBI, blood flow is disrupted in both impacted and non-impacted regions of the brain, creating a hypoxic environment that may influence ciliary gene and protein expression on ECs.

## Introduction

Traumatic brain injury (TBI) is a pressing public health problem impacting ∼2 million people and resulting in >60,000 deaths in the United States each year^1–3^. In addition to neuronal damage or dysfunction, TBI is also known to injure the cerebral blood vessels leading to hemorrhage, edema, and changes in cerebral blood flow (CBF)^4^. Depending on the severity of the injury and associated vascular sequelae, TBI can result in three phases of altered CBF. Initially, hypoperfusion or decreased blood flow occurs, and this is followed by hyperemic or increased blood flow, before transitioning to another period of decreased blood flow often caused by vasospasm^5^.

In the acute phase, within 72 hours of severe TBI (sTBI), CBF reduction and the prolongation of mean transit time of the cerebral circulation have been correlated with worse clinical outcomes^6^. Magnetic resonance imaging assessments showed that CBF alterations have been detected 3-6 months post-injury suggesting TBI results in long-term vascular dysfunction or remodeling^7^.

Altered CBF assessed 3-6 months post-injury also correlates with changes in executive function, which implies an important role of abnormal CBF in long-term morbidity^7^. Vascular remodeling changes after TBI include a reduction in the overall volume of cerebral blood vessels suggesting that TBI may result in chronic brain hypoperfusion^8^. Post-TBI cerebrovascular alterations including CBF changes are also observed in pre-clinical TBI animal models^9–13^. Collectively, these data support the hypothesis that TBI results in acute cerebrovascular damage that can evolve into a chronic condition that includes abnormal CBF. Thus, following sTBI, the underlying vascular pathology may persist and remain untreated as we are currently limited by a lack of knowledge on CBF-associated vascular cell specific changes.

To better understand the effect of TBI at the cellular level, the advent of genomic approaches such as single cell RNA sequencing (scRNASeq) has revealed novel insights into TBI pathology^14–21^. Specifically, studies using a non-penetrating closed-skull injury mild TBI model identified impact on non-vascular cell clusters in various parts of the brain^14, 15, 18, 20^, and immune response post-injury^14, 16^ . However, molecular underpinnings of the consequence of sTBI on the brain vasculature remains under-appreciated so far. Flow in the vasculature is sensed by a microtubule-based organelle called cilia, which protrude from the cell membrane into the vessel lumen. EC-cilia act as sensors of blood flow and metabolism and translate information about the environment to activate the endothelium^22–24^. Cilia are known to have important roles in several functions to maintain vascular health such as regulating vascular integrity, inducing the production of nitric oxide, and regulating cellular immune responses ^23, 25–27^. Previous work from our group showed that altered flow conditions affect brain EC cilia structures leading to damage and release of ciliary proteins^28^. To date, whether flow-associated changes post TBI influences EC-cilia is not known. We hypothesized that changes in CBF post sTBI will damage EC-cilia and influence genes expressed in ECs suggesting their activation in response to the injury. To test this hypothesis, we used a closed-skull mouse sTBI model to characterize the temporal changes in CBF and employed scRNAseq to detect ciliary gene changes specifically in ECs in the injured brain.

## Methods

### 1.1 Mice

All mouse experiments were approved by the Medical College of Wisconsin Institutional Animal Care and Use Committee. C57BL/6J mice were purchased from Jackson Laboratories (Bar Harbor, ME) and bred under specific-pathogen-free conditions at the MCW Biomedical Resource Center. A breeder pair of ARL13B-EGFP^tg^ mice were generously donated by David Clapham (Janelia Research Campus, Ashburn, VA, USA)^29^ to create a colony maintained by the Rarick laboratory. All ARL13B-EGFP^tg^ mice used were from the Rarick colony. All experiments were performed on 12- to 14-week-old mice. Animals used in this study were housed in a 12- hour light/12-hour dark cycle with free access to food (standard mouse chow) and water ad libitum. Animals were monitored by lab staff and animal facility staff, which included full-time veterinarians. For the end point tissue collection, mice were deeply anesthetized using inhaled isoflurane (4%), a thoracotomy was performed to access the heart to draw blood samples followed by trans-cardiac perfusion to flush and clear circulating blood from the organs with heparinized phosphate buffered saline (PBS).

### 1.2 Traumatic brain injury (TBI)

TBI was applied in age-matched (12- to 14-week-old) male C57BL/6J mice using the non-surgical, closed-skull modification of the controlled cortical impact model^30^. The closed-skull modification maintains the integrity of the skull and meninges allowing for appropriate changes to intracranial pressure and cerebrospinal fluid dynamics and avoiding complications that may occur with a surgical craniotomy including non-TBI related inflammation and cellular responses. A severe TBI with hemorrhage was created using a single impact with the following parameters: 5 mm diameter tip; 2 mm impact depth, 0.2 s dwell time, and 6 m/s velocity (Leica MyNeuroLab Impact One Stereotaxic Impactor, Leica Biosystems, Richmond, IL). To induce TBI, mice were anesthetized in an induction chamber using 4% inhaled isoflurane. Mice were then moved under the impact device with their head positioned on a bed of gauze perpendicular to the impact tip with isoflurane anesthesia delivered via nosecone maintained at 1.5%. This allowed free movement of the animal’s head in response to the impact. The impactor tip was positioned over the right parietal bone at ∼10° angle to achieve greater contact between the flat surface of the tip and the curved skull. Anesthesia was stopped immediately after impact limiting the total time of anesthesia exposure to less than 5 minutes for the procedure, including induction. Body temperature was maintained during the procedure and recovery periods by placing the mice on a heated recirculating water pad.

### 1.3 Laser speckle contrast imaging (LSCI)

LSCI was done to assess the impact of TBI on blood flow changes in the cerebral cortex. For LSCI, the animal was anesthetized using inhaled isoflurane (2.5%) delivered through a nose cone with 24% O2/76% N2. A midline incision in the scalp was made and the scalp retracted to expose the skull. The skull remained intact and covered with sterile saline to prevent drying. During imaging, the anesthesia was lowered to ∼1.5% to minimize cardiorespiratory depression and allow for robust cerebrovascular reactivity. After lowering the anesthesia, cerebral perfusion was monitored for at least 15 minutes to ensure values were stable then perfusion images were collected. Two equal sized regions of interest (ROI) were drawn to include the entire parietal cortex of each brain hemisphere (impacted side = ipsilateral hemisphere, non-impacted side = contralateral hemisphere). Data presented are the average perfusion value calculated for each ROI. Separate cohorts of mice were used to evaluate 1) the immediate cerebral perfusion changes during the first 2 hours after sTBI; 2) the acute changes in cerebral perfusion comparing measurements acquired pre-TBI, 30 minutes post-sTBI, and 7 days post-sTBI; and 3) the chronic effect on cerebral perfusion measurements acquired 28 days post-sTBI.

### 1.4 Flow cytometry

At the respective endpoints, 5 mm x 5 mm brain cortex samples were collected from the site of injury after the brains were perfused as described above. The area and the volume of tissue samples remained consistent within mice. Similar to the cortex samples harvested from the side of injury, specimens were also collected from the symmetrical site-matched un-injured contralateral side. Matching cortex samples from no-TBI control mice were also collected.

Cortex samples were finely dissected and digested in 5 ml buffer (1 mg/ml collagenase D, 100 µg/ml DNase, 5 mM CaCl2, papain suspension 60 U, 10% fetal bovine serum (FBS) in RPMI medium) in a dish for 30 minutes inside a shaker maintained at 80 RPM, 37°C. To remove any undigested fraction or tissue particles, samples were then passed through a 40 µm cell strainer (pluriSelect, cat# 43-50040-01). Myelin debris were then removed using specific magnetic beads (Miltenyi Biotech, cat# 130-096-733) per manufacturer’s recommended protocol. Single-cell suspensions were then washed twice with FACS buffer (1× PBS with 5% FBS and 0.1% NaN3) at 300g for 5 minutes and were subsequently incubated with Live/Dead fixable yellow dead cell stain (Thermo Fisher cat# L34968) as per manufacturer’s protocol, to exclude any dead cells, wherever applicable. Then, cells were fixed and permeabilized using Cytofix/Cytoperm buffer (BD, cat# 554722) and stained with the following antibodies to identify different blood-brain-barrier associated cell clusters (CD31+ cells as EC population, CD31-CD184+GFAP+ cells as astrocytes or glial cells) and cilia proteins: CD31-BUV396 (BD, cat# 740231), CD184-PerCP Cy5.5 (Biolegend, cat# 146509), purified monoclonal GFAP (Biolegend, cat# 644702) and purified polyclonal ARL13B (Proteintech, cat# 17711-I-AP). Suitable secondary reagents were used to detect the respective primary antibodies. Primary antibodies were diluted 1:100, and secondary antibodies were diluted 1:500. BD perm wash buffer (cat# 554723) was used for antibody dilutions and washing. Primary antibodies were incubated for 45 minutes and secondaries for 30 minutes at 4°C. Suitable secondary antibody controls were included. After the completion of staining, cells were resuspended in FACS buffer. Stained cells were run on a flow cytometer (Becton Dickinson LSRFortessa). Sample acquisition was done using Becton Dickinson’s FACSDiva software with subsequent analysis on FlowJo software.

### 1.5 Tracking cilia protein in blood

ARL13B-EGFP^tg^ mice were subjected to TBI as described above. 50 µl of blood was collected in heparinized tubes from each mouse pre- and post-TBI from the jugular vein. Blood cells were fixed in 4% paraformaldehyde and labeled with antibodies to detect the cilia protein ARL13B as described above. After staining, cells were resuspended in FACS buffer and run in the flow cytometer as described above. The transgenic EGFP signal was detected to validate the ARL13B signal as detected by the antibody. At least 50,000 blood cells were run per sample.

### 1.6 Single cell RNA sequencing

Single cell suspensions were prepared from brain cortex samples post-injury (Days 1, 7, and 28) and from no-TBI control mice using the cell isolation procedure described for flow cytometry. 5 mm x 5 mm cortex specimens were collected specifically from the site of injury and site-matched cortex from un-injured control mice. After removing myelin debris, single cell isolations were fixed and processed for library construction as per manufacturer’s (10x Genomics) protocol. For multiplexing, samples were probed using the single cell fixed RNA transcriptome probe kit. For a single library construction, 4 samples from each time point were multiplexed by the dual index plate reagent following the preparation of gel emulsion using the 10x Genomics Chromium X instrument. For sequencing, 4 libraries were multiplexed. Each library is identifiable by a specific set of i5 and i7 primers as included in the dual index plate. Following the construction as recommended by the manufacturer, cDNA library products were validated with TapeStation (Agilent Technologies), by using High Sensitivity D5000 ScreenTape and analyzing with TapeStation analysis software version 3.1. The Chromium X (10x Genomics) generated barcoded gel-bead emulsions from post-fixed single cell, and Illumina-compatible library preps were subsequently sequenced on Illumina High Seq-2500 platform. We sequenced ∼10,000 cells per sample. Feature-barcode matrix was generated using the Cell Ranger 7.1.0 (10x Genomics). Sequenced data were aggregated, and reads were aligned by the Cell Ranger. The following quality control steps were conducted: 1. Cells that have unique feature counts over 3500 or less than 200 were excluded for further analysis; and 2. Cells that have over 5% of unique molecular identifiers were derived from the mitochondrial genome were excluded for further analysis. Seurat package was used for data normalization, principal component analysis, Uniform Manifold Approximation and Projection (UMAP), clustering analysis and differential expression analysis. Genes were considered significantly differentially expressed at a false discovery rate of 5%. Gene set enrichment analyses (GSEA) were performed to identify important pathways. Based on FindMarkers function from Seurat, we used gseKEGG function in ClusterProfile Package^31^ for GSEA. Library org.Mm.eg.db was used for gene ID mapping. We used dotplot and gseaplot2 for further visualizing the GSEA results.

### 1.7 Cell culture

Primary human brain microvascular endothelial cells (HBMVECs, Cell Systems Corporation cat# ACBRI 376) were cultured in 10 cm culture dishes or 6-well tissue culture plates on glass coverslips and maintained at 37°C in a 5% CO2 incubator in endothelial cell complete medium (Promocell, cat# C22010). Cultured HBMVECs at 90% confluency were either maintained under normoxic condition (room air) or subjected to hypoxia (2% oxygen) in the cell culture incubator for 24 hours and then collected for analysis. Hypoxic conditions inside of the incubator were maintained using the ProOx 110 Compact O2 Controller (BioSpherix, Ltd). A separate group of hypoxia treated HBMVECs were returned to normoxic conditions for an additional 24 hours and then collected. To validate that hypoxic conditions were achieved, we measured HIF1-alpha protein changes using flow cytometry. We also used a fluorogenic hypoxia indicator dye loaded into the cells cultured on cover slips as per manufacturer’s recommended protocol (Image-iT™ Green Hypoxia Reagent, ThermoFisher cat#I14834). HBMVECs cultured in 10 cm dishes were processed and analyzed as described above using flow cytometry to detect protein changes. HBMVECs cultured in six-well plates on coverslips were washed twice with 1X PBS (Gibco, Cat#10010023) and subsequently fixed with 4% PFA (Electron Microscopy Sciences, Cat#15710) for 15 minutes and imaged using a Zeiss confocal microscope at a magnification of ×63 to detect the green, fluorescent hypoxia reagent. Images of HBMVECs were captured from areas of similar confluence on each cover slip.

### 1.9 Statistics

Data were presented as mean and standard deviation (SD), or standard error of the mean (SEM) as noted. Changes in cerebral perfusion measured using LSCI were analyzed using repeated measures two-way ANOVA and each time point post impact was compared to the pre-TBI value. A two-tailed two-sample t test or one-way ANOVA was performed to compare differences between groups as appropriate based on the number of comparisons. The differences of % total Arl13b+ blood cells between pre- and post-TBI were evaluated by a paired t test. Kruskal-Wallis Test was used where parametric assumptions were not satisfied. Dunnett’s test, Sidak’s, or Tukey’s test was used to adjust for multiple comparisons. P < 0.05 was considered statistically significant. Statistical analysis was performed using SAS V9.4 (SAS Institute Inc.) and GraphPad Prism software (version 10.2.0).

## Results

### Cerebral blood flow is altered post TBI

We monitored blood flow changes in the brain cortex using LSCI before and after sTBI. We observed an immediate decrease in CBF in response to the impact that transiently recovers before decreasing to a more sustained ischemia lasting for at least two hours **(Fig 1A, Supp Fig 1)**. Interestingly, blood flow was compromised not only in the impacted side of the brain (ipsilateral), but also in the non-impacted hemisphere (contralateral). By day 7 post-sTBI, the cerebral perfusion tended to be higher as compared to pre-TBI **(Fig 1B, Supp Fig 2)** suggesting a state of hyperemic blood flow, although this varied by animal due to apparent differences in sustained injury severity. Chronically altered cerebral perfusion patterns continued to be present at day 28 post-sTBI where we observed regions of reduced blood flow that corresponded to the impact induced lesion **(Supp Fig 3)**. Collectively, our data suggests that sTBI causes blood flow disturbances that transition as the injury evolves which can remain unresolved even after 4-weeks post-injury.

**Fig 1:**
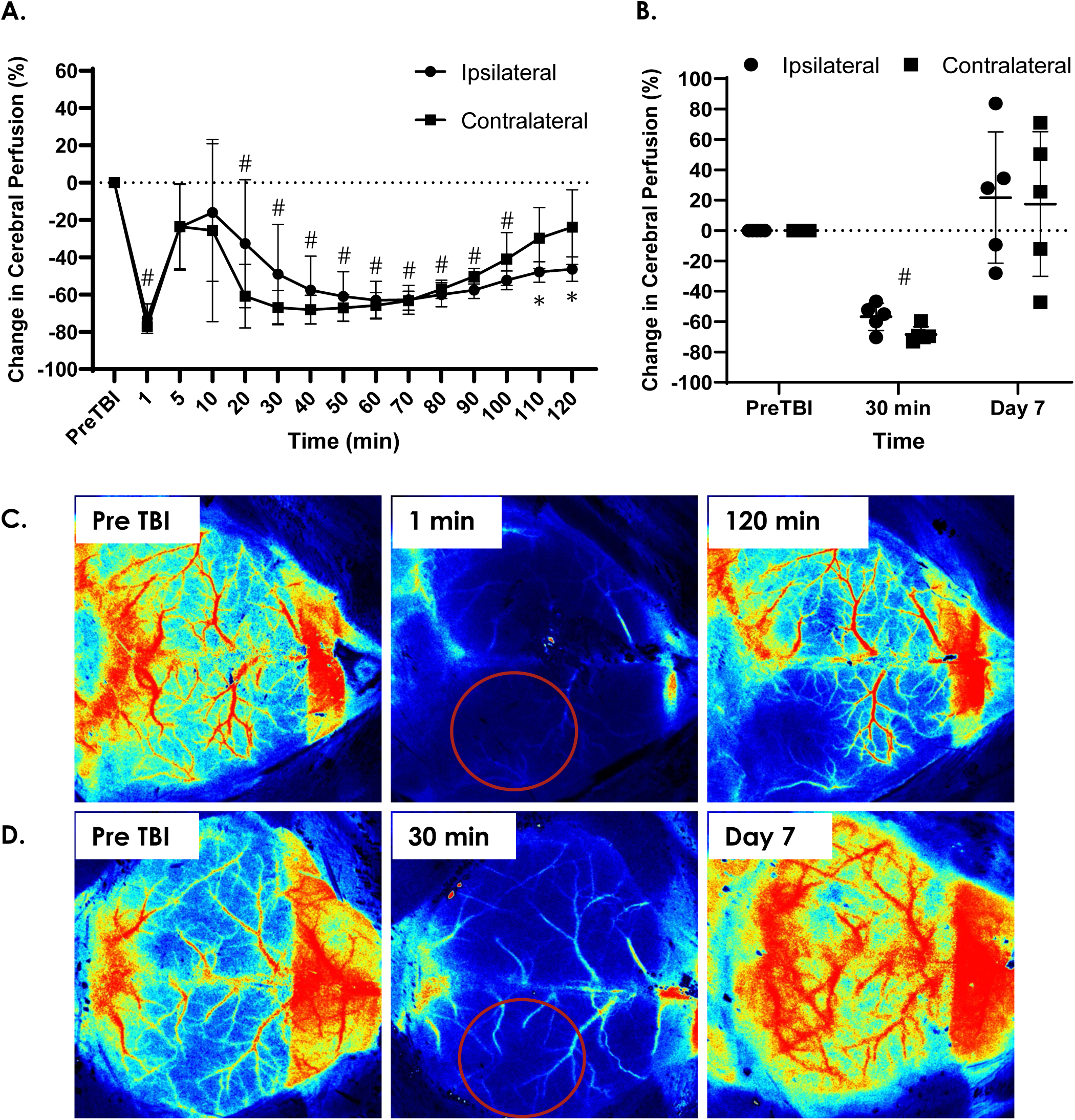
T**B**I **causes altered blood flow in brain.** LSCI was used to measure cerebral perfusion before and after sTBI. Percent change in cerebral perfusion before and up to 120 minutes after sTBI (n=4 mice, # = p<0.05 for both ipsilateral and contralateral hemispheres compared to each region’s pre-TBI value, * = p<0.05 for ipsilateral hemisphere compared to its pre-TBI value (A). Percent change in cerebral perfusion before, 30 minutes post impact, and seven days after sTBI (n=5 mice, # = p<0.05 for both ipsilateral and contralateral hemispheres compared to each region’s pre-TBI value (B). Representative LSCI images for data shown in panel A (C) and panel B (D). Site of impact denoted by red circles. Data are mean ± SD. Repeated measures Two-way ANOVA was used to determine the change in perfusion over time compared to pre-TBI values. Dunnett’s test was used to correct for multiple comparisons.

**Fig 2:**
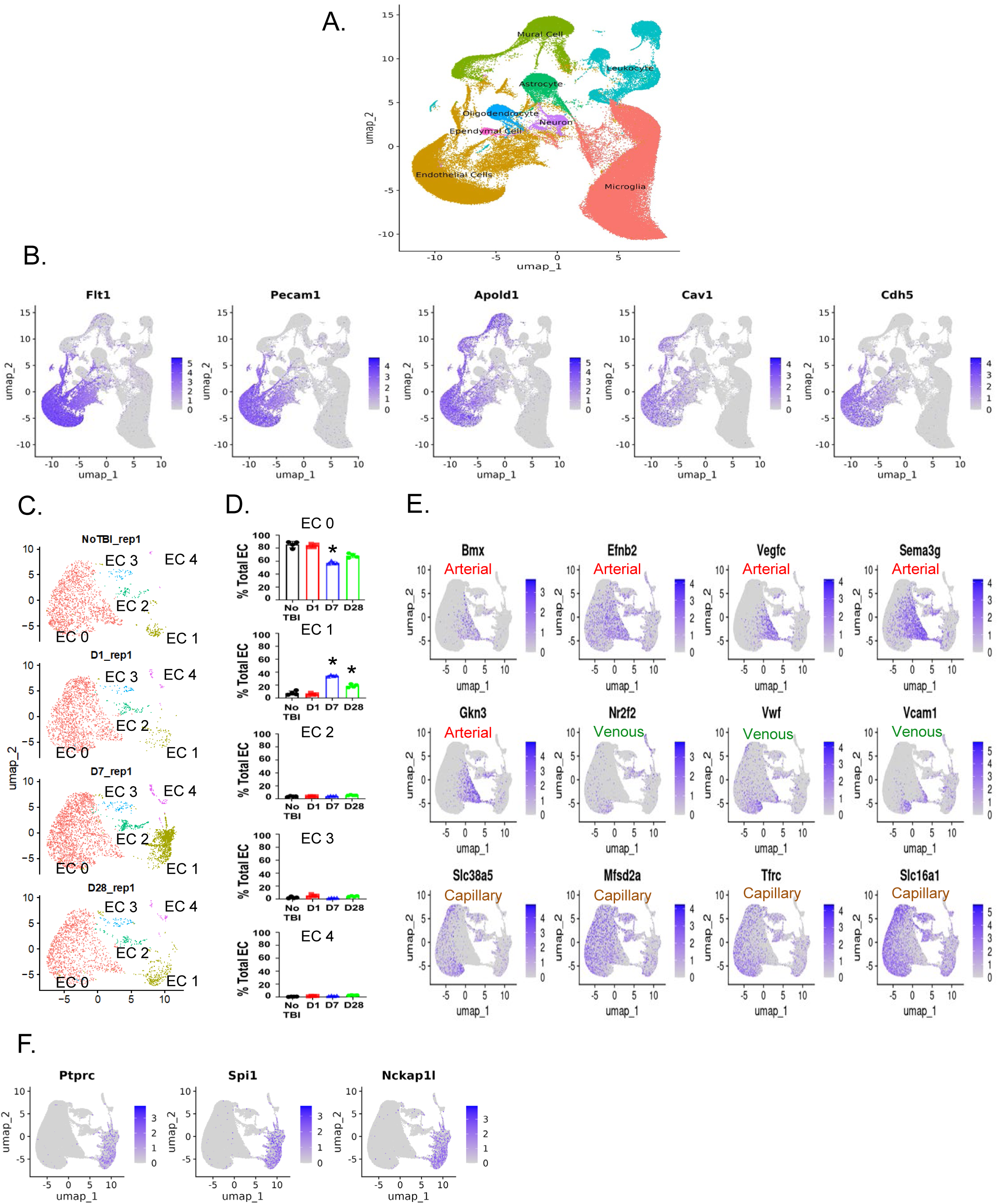
s**c**RNAseq **to identify brain-derived EC clusters post-TBI.** Representative UMAP demonstrates how scRNAseq identified brain resident cell clusters including EC (A). Feature plots show EC-specific gene enrichment in UMAP-derived cell clusters (B). Representative UMAP plots demonstrate the resolution of 5 EC sub-clusters (C). Quantification of temporal changes in representation of EC subclusters (D). Representative feature plots to show zonation-specific arterial, venous, and capillary gene enrichments within EC sub-cluster 0 (E). Representative feature plots to show hematopoietic gene signature of EC sub-cluster 1 (F). For each time point, n=4. Data are mean ± SEM. * p<0.05 vs. No TBI. Kruskal-Wallis and ANOVA tests were performed respectively for EC cluster 0 and 1. Dunn’s test and Turkey’s test were performed for multiple comparison adjustment respectively for EC cluster 0 and 1.

**Fig 3:**
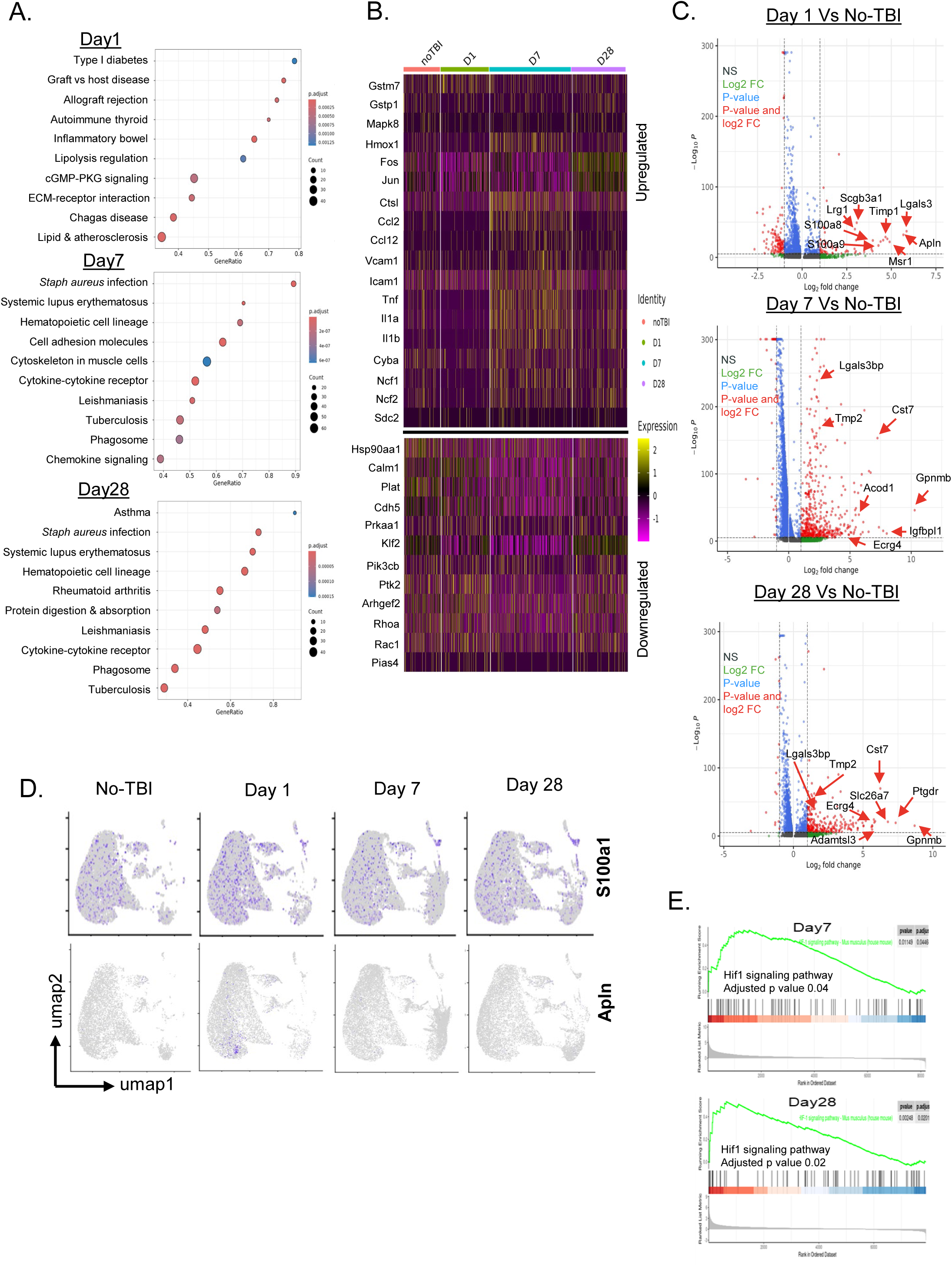
s**c**RNAseq **to identify molecular pathways in brain-derived EC post-TBI.** Pathway analysis on EC cluster at day 1, 7 and 28 post-TBI (A). Heatmaps show the list of genes from the shear stress pathway that were upregulated (top panel) or downregulated (bottom panel) in EC cluster post-TBI (B). Volcano plots with top significantly differentially expressing genes in EC cluster respectively at day 1, 7 and 28 post-TBI (C). Genes from ischemia or hypoxia pathways are highlighted. Representative feature plots demonstrating the enrichment of additional ischemia-associated genes in EC cluster (D). Gene set enrichment analysis of hypoxia pathway in EC cluster at day 7 and 28 post-TBI (E). For each time point, n=4.

### Altered dynamics of EC sub-clusters post-TBI

To investigate the cellular changes at the site of injury where the altered CBF was most pronounced, we isolated brain tissue from the impacted cortex and subjected them to scRNAseq. Data from this effort was processed further at the bioinformatic level, and subsequently analyzed for the transcript expression changes specifically in brain ECs, the first line of cells directly in contact with altered blood flow and known to be flow responsive. We observed a single electropherogram peak with a size of 260bp for each study time point validating the optimal quality of our RNA library preparations according to the manufacturer’s protocol **(Supp Fig 4)**. We then utilized Seurat-based unsupervised UMAP algorithm that reduces a high-dimensional RNAseq data set to a lower dimension with minimal loss of the original data. Seurat-based unsupervised UMAP algorithm identified cells belonging to the EC cluster **(Fig 2A)**. This cluster showed expressions of common EC-specific genes such as *Flt1, Pecam1, Apold1, Cav1 and Cdh5* **(Fig 2B)**. We then utilized a higher order UMAP algorithm to identify 5 sub-clusters of ECs **(Fig 2C)**. EC sub- cluster 0 (red colored cluster in **Fig 2C)** remained the most prominent sub-cluster that constitutes ∼60 – 90% of total EC cluster **(Fig 2D)**. The proportion of this sub-cluster remains the same as baseline 1 day after the injury, was significantly reduced by ∼30% at day 7, and then increased towards baseline levels again at day 28 post-TBI. EC sub-cluster 1 **(Fig. 2C**, green in color) remains the second most prominent EC-subcluster that constitutes <10% of total EC cluster until day 1 post-injury, peaks to ∼35% at day 7 and remains at >20% even after day 28 post-TBI **(Fig 2D)**. Each of three other EC-subclusters constitutes <5% of total EC and their respective proportions did not change over time post-injury **(Fig 2D)**. EC subcluster 0, the most abundant subcluster showed enrichment for genes associated to zonation-specific arterial (*Bmx, Efnb2, Vegfc, Sema3g, Gkn3*), venous (*Nr2f2, Vwf, Vcam1*), and microvascular signatures (*Slc38a5, Mfsd2a, Tfrc, Slc16a1*) **(Fig 2E)**^32^. The top 10 differentially expressed genes in EC sub-cluster 1 are listed in **Table 1**. Interestingly, of these, 3 genes (*Ptprc, Spi1, Nckap1l*) were hematopoietic cell-associated **(Fig 2F)**. This endothelial RNA-seq data analysis in sTBI brain suggests that a unique population of ECs at or before day 7 are emerging as the TBI pathology progresses over time.

**Fig 4:**
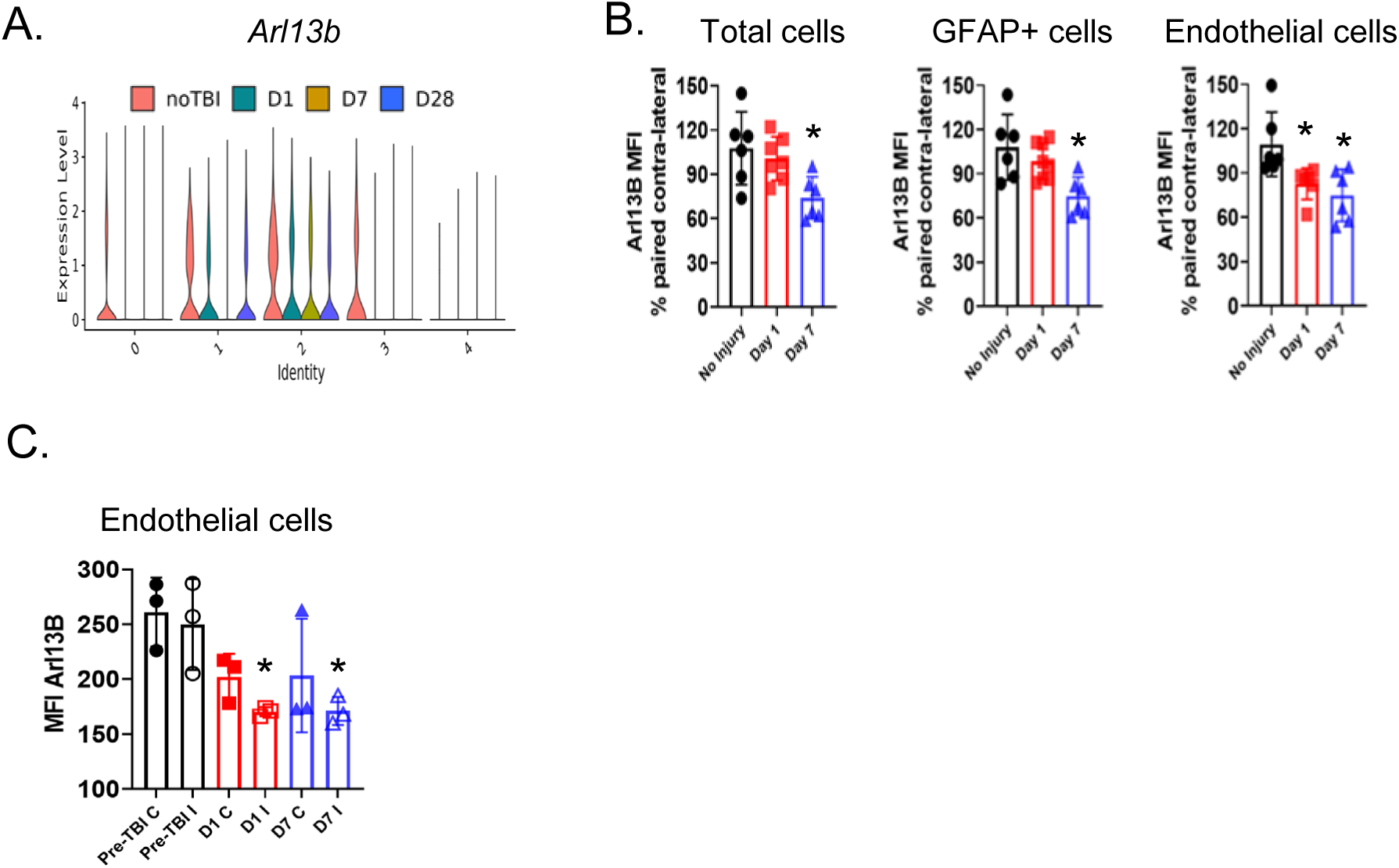
E**C**s **decrease expression of ciliary markers post-TBI.** scRNAseq identifies the loss of *Arl13b* gene expression from EC clusters (A). Expression of ARL13B protein was quantified by flow cytometry from total cells, GFAP+ cells, or ECs derived from brain cortex. For any given post-TBI time point or uninjured control (no-TBI), the ARL13B expression in single cells from ipsilateral side was normalized against single cells from site-matched paired contralateral side. Each group of mice were euthanized on separate days at the indicated endpoints (B). In an additional experiment, expression of ARL13B protein was quantified by flow cytometry from endothelial cells derived from brain cortex. Here, each group of mice were injured on separate days to coincide all endpoints on a same day. Thus, all groups of mice were euthanized the same day but with different endpoints (C). For panel A, n=4. For panel B, n=6. For panel C, n=3. Data are mean ± SEM. * p<0.05 vs. control (Pre-TBI or No Injury). For statistical analysis, ANOVA or Kruskal-Wallis was performed; Tukey’s test or Dunn’s was used to adjust for multiple comparisons.

### Enrichment of vascular pathology and inflammatory pathways in ECs post-TBI

To further understand the local endothelial response in the brain at the site of the injury, we investigated the molecular pathways that were dysregulated in brain EC cluster post sTBI. We analyzed top differentially expressed genes to deduce specific molecular pathways that were up or downregulated post injury. Prominent altered vascular function-related pathways included lipid and atherosclerosis-related signaling pathways at day 1 as identified by GSEA **(Fig 3A)**. In addition, we observed several generic inflammatory pathways such as cell adhesions, chemokine signaling, and cytokine-cytokine receptor interactions were also upregulated at day 7 and 28 post-sTBI **(Fig 3A)**. TBI also resulted in upregulation of metabolism-associated pathways such as protein digestion and diabetes mellitus **(Fig 3A)**. Consistent to the expansion of EC subcluster 1, we also observed upregulation of hematopoietic cell lineage pathway at day 7 and 28 post-injury **(Fig 3A)**. These data collectively suggest that post injury, brain ECs are exposed to a metabolically altered inflammatory environment post-TBI. We further analyzed top 500 most differentially expressed upregulated genes in EC cluster post-TBI. Since we observed significantly altered CBF post-injury, we further analyzed flow-associated shear stress responsive gene sets in the EC cluster at any time point post-injury **(Fig 3B)**. Interestingly in this pathway, we observed upregulation of chemokine genes such as *Ccl2, Ccl12*; upregulation of adhesion molecule associated genes such as *Vcam1, Icam1*; upregulation of inflammatory cytokine genes such as *Tnf, Il1a, Il1b* at day 7 post-injury **(Fig 3B)**. The expression of inflammatory cytokine genes remained elevated even at day 28 post-injury. *Cdh5*, a gene corresponding to endothelial adherens junction protein cadherin 5, was found to be downregulated at day 7 post-injury. Interestingly, the expression of physiological shear-responsive gene *Klf2* was downregulated as early as day 1 and was not rescued even at day 7 post-injury **(Fig 3B)**. Collectively, the data suggests that exposure to altered blood flow conditions post-injury dysregulates the shear force-associated transcriptional programming in brain ECs.

### Enrichment of ischemic and hypoxic pathway in ECs post-TBI

To further characterize the transcriptomic signature of the brain ECs post-sTBI, volcano plots were generated with all top differentially expressed genes at the respective post-sTBI time points **(Fig 3C)**. Interestingly at day 1, among top differentially upregulated genes, we found at least 10 genes such as *Scgb3a1, s100a1, s100a8, s100a9, Apln, Lrg1*, *Chst*, *Timp1*, *Lgals3*, and *Msr1* which are implicated in ischemia or anti-ischemia host tissue responses **(Fig 3C and Table 2)**. This suggests that the reduced CBF that we observed acutely post-sTBI **(Fig 1A)**, is causing an ischemic or low oxygen condition in the brain, that affects ECs. Additionally, *s100a1*, a known acute ischemia- biomarker^33^ is enriched throughout all zones in EC subcluster 0 while ischemia-associated gene *Apln* is found upregulated exclusively in the venous zone of EC subcluster 0 **(Fig 3D)**. We also checked other s100 family genes, but none were found enriched in EC clusters **(Supp Fig 5).** The ischemic pathway remains active even at post-TBI day 7 and 28 as we observed upregulated expression of genes such as G*pnmb, Lgals3bp, Cst7, Timp2, CD72, Igfbpl1, Acod1, Ptgdr, Slc26a7, Adamtsl3* and *Ecrg4* **(Fig 3C)** which are implicated towards ischemia, ischemic stroke, or post-ischemic brain injury **(Table 2)**. Interestingly, at later time points post-sTBI (day 7 and 28), we observed significantly upregulated genes associated to hypoxia **(Table 2)** namely *Gpnmb, Timp2, Slc4a10* and *Ecrg4* **(Fig 3C)**. Gene set enrichment analysis revealed that the hypoxic pathway is indeed strongly enriched in ECs at day 7 and 28 **(Fig 3E)**, but not at day 1. The heatmap, volcano plot and GSEA enrichment analyses collectively suggest an association wherein altered CBF at day 1 creates an underlying ischemic environment that persists until day 28, and in between days 1-7, a new EC cluster emerges that is concomitant with the injury-induced state of hypoxia.

**Fig 5:**
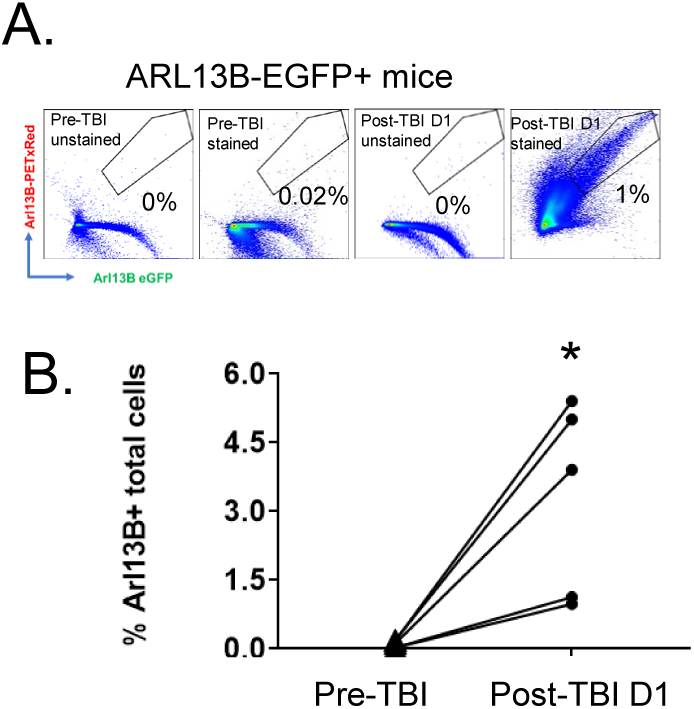
C**i**lia **protein ARL13B is detected in blood post-TBI.** ARL13B-EGFP mice were used to track and validate ARL13B in blood (A). Blood was drawn before and after the injury. Blood cells were immediately fixed, stained for ARL13B, and run in flow cytometer. Transgenic EGFP signal validates antibody bound ARL13B signal. Representative dot plots show the detection of ARL13B from injured mice, along with no-antibody (unstained) control samples (A). Percent total ARL13B+ blood cells were quantified at day 1 post injury as well as from paired pre-injury samples (B). n=5. * p<0.05 vs. Pre-TBI. For statistical analysis, paired t test was performed.

### Brain ECs lose cilia protein ARL13B post-TBI

As we observed persistent altered CBF in brain post-injury, and in accordance with our previous observation that altered flow causes loss of cilia^28^, we further investigated any change in cilia-specific markers at the gene and protein levels. Most cilia genes were prominently expressed in ependymal cells and at very low levels in EC clusters **(Supp Fig 6A-B)**. Cilia gene *Arl13b* expression was uniquely compromised in EC sub-clusters 0 and 3 at day 1 post-TBI and such loss was never recovered **(Fig 4A)**. *Arl13b* expression was uniquely low in EC sub-cluster 1 at day 7 post-TBI, when the cluster was observed to be significantly expanding **(Fig 4A)**. Progressive loss of *Arl13b* gene expression over time was also noted in EC sub-cluster 2 **(Fig 4A)**. Six cilia genes, namely *Pafah1b1, Tuba1a, Gsk3b, Ctnnb1, Dpysl2* and *Pcm1*, were the other top candidates that were detectable in ECs **(Supp Fig 6C)**. Of these, *Pafah1b1* expression was mostly detectable in no-TBI group, and this was consistent to the expression pattern of *Arl13b* **(Supp Fig 6C)**.

**Fig 6:**
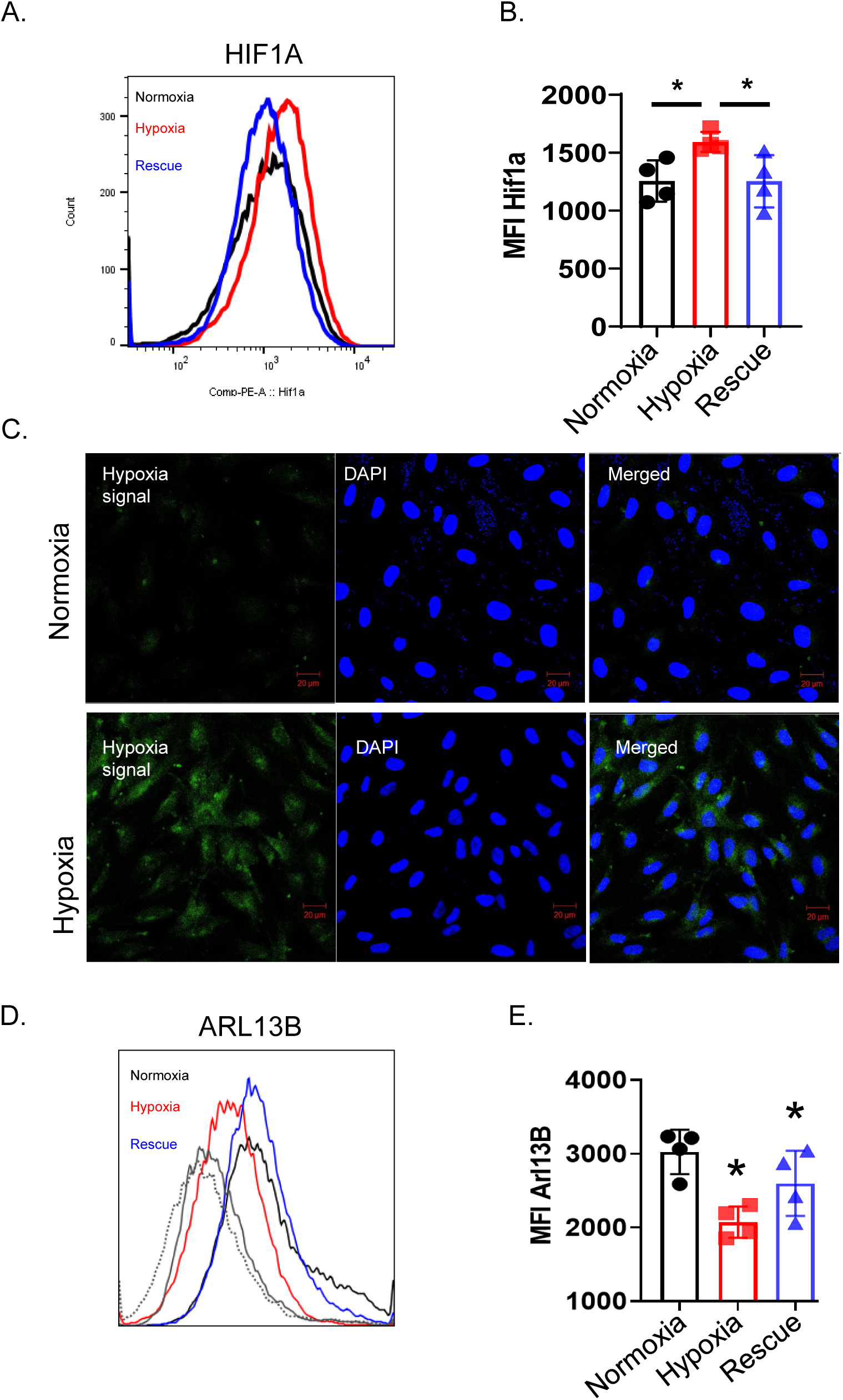
H**y**poxia **causes loss of cilia in EC in vitro.** Human brain microvascular ECs were cultured under normoxia or hypoxia (2% oxygen saturation) conditions for 24 hours. A third group of cells were cultured in hypoxia for 24 hours followed by normoxia for an additional 24 hours (Rescue). Cells were subsequently run in the flow cytometer to measure changes in HIF1A and ARL13B protein expression. Representative histogram showing the expression of HIF1A in different experimental groups and the corresponding quantification in terms of median fluorescence intensity (MFI) (A-B). In experimental duplicates, cultured cells were stained with hypoxia reagent and green hypoxia positive signal was detected by immunofluorescence to validate experimental hypoxia *in vitro* (C). Total cellular ARL13B ciliary protein was quantified by flow cytometry in experimental duplicates. Representative histogram showing the expression of ARL13B in different experimental groups. Unstained cells (gray solid) and secondary antibody control (gray dotted) were also included (D). Corresponding quantification of ARL13B MFI (E). n=4. For panel B and E, n=4. Data are mean ± SEM. * p<0.05. For statistical analysis of hypoxia validation (B) ANOVA was performed; Tukey’s test was used to adjust for multiple comparisons. For statistical analysis of Arl13b MFI (E), two sample t test was performed.

**Fig 7:**
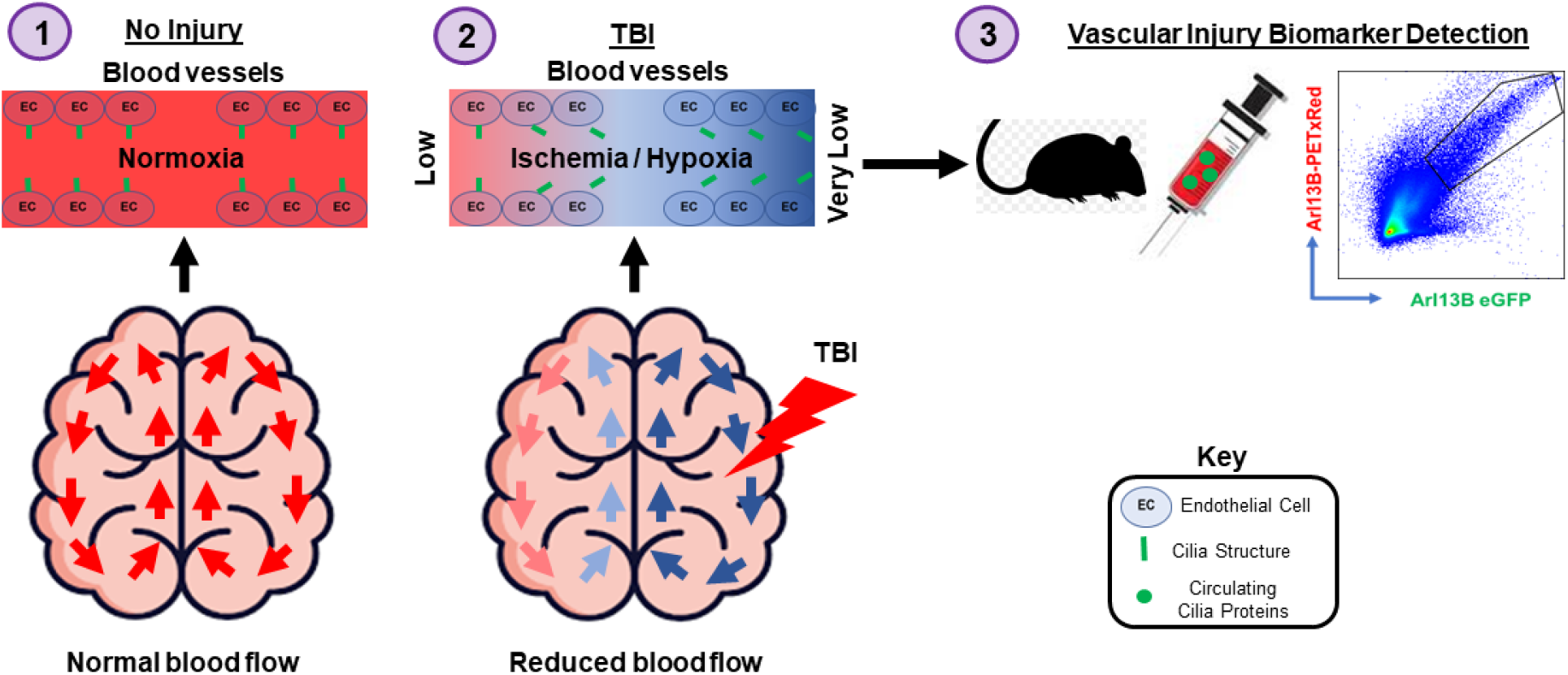
S**c**hematics **of the concept.** Normal blood flow in brain under the normoxic condition pre-TBI (1). Following TBI, altered blood flow condition is established. Reduced blood flow contributes towards ischemic and hypoxic condition in brain endothelium (2). Brain ECs lose cilia due to altered flow or metabolic condition, and the cilia protein is trackable from blood (3).

We also used flow cytometry to quantify changes in cilia protein expression in single cells isolated from injured ipsilateral and paired un-injured contralateral cortex samples at day 1 and 7 post-injury. We did not see any significant change in the expressions of cilia protein ARL13B in total cells harvested from the injured ipsilateral side, normalized to uninjured contralateral side at day 1, but the expression of the protein was significantly reduced in ipsilateral ECs at day 7 post- sTBI **(Fig 4B)**. We observed a similar trend of ARL13B expression in astrocytes, detected as GFAP+ cells **(Fig 4B)**. Interestingly at day 1 post-sTBI, expression of ARL13B was significantly reduced in ipsilateral ECs, as normalized to contralateral ECs, and remained so even at day 7 post-sTBI **(Fig 4B)**. Because cerebral perfusion was reduced in the non-impacted contralateral hemisphere, we conducted an additional experiment to compare cilia protein expression within each brain hemisphere across all study timepoints. In this experimental duplicate, we injured different groups of mice on different days to collect and analyze all study endpoints on the same day. Compared to no-TBI cortex-derived ECs, we observed significantly reduced ARL13B in ECs harvested from the impacted (ipsilateral) cortex at day 1 **(Fig 4C)**. Interestingly, the cilia protein ARL13B was not only reduced in the ECs harvested from the side of injury, but also reduced in ECs from the contralateral non-impacted side **(Fig 4C)** suggesting a diffuse injury throughout the brain cortex that may have resulted from the global alterations in CBF observed after sTBI.

### Cilia proteins are tracked in blood post-TBI

We hypothesized that cilia proteins that are lost from the brain ECs post-sTBI, are either found in blood or on blood cells. Thus, we assayed the abundance of cilia proteins in the blood collected from ARL13B-EGFP transgenic mice^29^.

Samples from the transgenic mice were additionally labeled with a commercially available anti-ARL13B antibody (PE-Texas red signal), and subsequently EGFP+PE-Texas red+ signal was identified to confirm any presence of ARL13B protein in the blood **(Fig 5A)**. While ARL13B was barely detectable in blood pre-TBI, the protein was abundant and associated to >5% of total blood cells at day 1 post-sTBI **(Fig 5B)**.

### Hypoxia causes the loss of cilia proteins in brain ECs

We further investigated whether hypoxia could contribute directly to the endothelial loss of cilia proteins. In our *in vitro* culture system using HBMVECs, we established hypoxia (2% O2), and validated the hypoxic condition by measuring changes in HIF1A protein expression using flow cytometry **(Fig 6A-B)** and using a fluorogenic compound that begins to fluoresce when atmospheric oxygen levels are below 5% **(Fig 6C)**. We detected cellular ARL13B protein content under normoxia or hypoxia and quantified total ARL13B by flow cytometry **(Fig 6D-E)**. Compared to normoxia, the cellular expression of ciliary protein ARL13B was significantly reduced under hypoxia. Collectively our data suggests that brain ECs lose cilia proteins not only due to the impact of the injury, but also under the altered metabolic environment as originated from the reduced CBF post-sTBI.

## Discussion

In the current study, we tracked severely compromised CBF not only in the impacted cortex, but also in the non-impacted, contralateral cortex. Our study demonstrates that altered CBF is established very quickly, globally affects the whole brain, and focal flow disturbances remain even at day 28 post-sTBI. Our scRNAseq data also revealed that as a pathological outcome of compromised CBF, an ischemic-hypoxic cellular environment is established in the endothelial cluster at least at the site of injury, and it appears to influence cellular signaling until day 28 post-sTBI. Using scRNAseq approach, we also identified five EC-subclusters that demonstrated different temporal gene expression patterns that may provide insight into the evolving vascular pathology after TBI. We have also identified that EC cilia, blood flow responsive cellular membrane structures, are directly influenced by low oxygen environments, which may be established post-injury due to altered CBF. Our study suggests that the abundance of cilia-derived protein in blood could prognosticate vascular injury or stress associated to altered CBF at least at day 1 post-TBI, which is consistent with our previous work identifying cilia as a blood flow associated indicator of vascular health^28^.

Consistent with previous reports^34, 35^, we observed post-TBI cerebral hypoperfusion not only in the impacted hemisphere, but also in the non-impacted contralateral hemisphere. The onset of cerebral hypoperfusion was both immediate and sustained for hours after impact. After the initial period of ischemia, we detected hyperemic blood flows in three out of five mice seven days after sTBI suggesting a secondary phase conducive to reperfusion injury. Due to the timing of LSCI measurements and variability in the severity of injury as suggested by the visual appearance of the impact lesions **(Supp Fig 2)**, the transition from ischemic to hyperemic blood flows in two of the mice likely occurred at different days as the time course of expected blood flow changes is likely due to the initial hemorrhagic insult^36^. Perfused brains collected at post injury day 7 and 28 demonstrate red blood cell breakdown products remaining within the parenchyma **(Supp Fig 2- 3)**. Severity of the injury and clinical outcomes are known to be related to the volume of hemorrhaged blood and the ongoing presence of residual blood products that remain after trauma with a hemorrhagic contusive injury component^37^. After hemorrhage, extravasated red blood cells lyse releasing multiple damaging factors such as hemoglobin, heme, and iron that need to be cleared to prevent ongoing secondary brain injury, including permeability of the BBB and development of cerebral vasospasm^36, 38, 39^. Collectively our data suggest that CBF disturbances are an acute and chronic vascular complication that cells throughout the entire brain deal with after TBI with hemorrhagic injury.

Our data indicates that sTBI and reduced CBF result in dysregulation of multiple immune and metabolic signaling pathways within 24 hours post-injury. Most of the top 10 dysregulated pathways are immune related, and many of the involved genes are also associated with vascular pathology specific to ischemia-reperfusion, hypoxia, or shear stress pathways **(Fig 3A-B)**.

Indeed, ischemia in TBI patients is known to correlate with poor clinical outcome^40^. Concurrent ischemia and impaired CBF are indicative of dysregulated autoregulation of cerebral circulation, a condition associated with poor neurological outcome in TBI patients^41, 42^. Interestingly the ischemia associated genes *S100a1* and *Apln*^43–46^ were highly expressed respectively in the microvessel and venous region specifically at day 1 post-TBI **(Fig 3D)** which suggests that specific blood vessel types are likely affected immediately following the injury or reactive to the altered metabolic environment established post-injury. The ‘low flow’ condition may result in metabolic deprivation such as lack of oxygen (hypoxia) at least for the brain endothelium.

Consistent to this notion, we observed upregulated hypoxia pathway which seems prominent in brain EC clusters at day 7 onwards. The continued enrichment of hypoxic pathway in brain EC clusters even at day 28 suggests that the consequence of initial post-injury vascular insult is long-term. Although previous work on mild TBI reported the enriched hypoxia pathway at day 1 post-injury, we did not observe this pathway activated early^14^. As the hypoxic environment promotes angiogenesis^47^, it is possible that the activation of hypoxic pathway facilitates post-injury vascular repair processes that might start earlier post mild injury, as opposed to later in severe injury. In addition, it is not clear why the EC clusters retained hypoxic gene signatures **(Fig 3C and E)** especially when the CBF demonstrates a hyperemic pattern. This question will require further examination; however, we did observe upregulation of several gene sets relevant to reperfusion injury including upregulation of shear-stress responsive and inflammatory cytokine genes as well as down regulation of the endothelial adherens junction gene cadherin 5. Our observation suggests that inflammation and cerebral metabolic disturbances persist during the hyperemic blood flow phase, possibly contributing to the long-term delay in returning to homeostasis.

Post-TBI, we observed the enrichment of hypoxia-associated gene expressions in the microvessel and venous EC clusters, which are associated with the blood brain barrier (BBB) and previously reported to be critical for tissue preservation post-injury^32, 48, 49^. Disruption of the BBB is a very early event post-TBI, that may persist long-term^50^. Thus, our data suggests that TBI impaired vascular homeostasis at least within the BBB regions. In support of this notion, we also observed upregulated expression of venous (*Vwf*) and capillary (*Tfrc*) genes along with upregulated BBB- preserving *Timp1* and *Timp2* gene expressions^51, 52^ in EC cluster at day 1 post-injury. This suggests a compensatory response from BBB-derived ECs following known structural and functional disruption at such barrier region post-TBI. Additionally, as previously reported, upregulated *Timp2* expression at day 7 is likely a compensatory response against post-TBI inflammation and hypoxia^52^.

Consistent to the altered CBF measured in the non-impacted contralateral side, we observed lower expression of cilia protein ARL13B in ECs isolated from the contralateral side post-TBI as compared to that of the no-TBI controls **(Fig 4C)**. This supports our hypothesis that altered flow condition causes the loss of endothelial cilia or cilia protein. Cilia is reported to be critical in maintaining the vascular integrity of the brain^25, 53^. Thus, the loss of cilia proteins from brain ECs at day 7 post-TBI implies that the structural integrity of brain vessels remains compromised.

Furthermore, we were able to detect increased concentrations of ARL13B protein in blood samples from mice one day after injury. These data collectively suggest that cilia proteins are released from brain ECs post TBI and may act as a biomarker of vascular injury. While several vascular biomarker candidates such as thrombomodulin, angiopoietin 2, von Willebrand factor and P-selectin have been previously associated with severe TBI^54^, currently only non-vascular proteins such as GFAP and UCHL1 have been cleared by the FDA for use as TBI biomarkers^55^. Our current work along with that of others suggests that a more complete characterization of TBI would be gained by using additional cellular biomarkers related to multiple aspects of the injury process such as vascular, inflammation, and metabolism.

In summary, our current study identifies that the loss of cilia is associated with TBI-induced altered cerebral perfusion. We observed post-TBI cerebral hypoperfusion even after the reported window of peak reparative angiogenic process^48^. This demands further investigation on bidirectional relationship between the CBF and the acutely injured vasculature, to better understand the post-TBI brain vascular homeostasis restoration process.

### Data Availability

Sequencing data for scRNA-seq is available on NIH Gene Expression Omnibus (https://www.ncbi.nlm.nih.gov/geo) with accession GSE271769.

### Sources of Funding

KRR, AG, AYP, RR, SMN, SPP, RZ were all partly supported by funds from HL154254 grant. This grant is part of the Trans-Agency Blood-Brain Interface program sponsored by NIH and the Department of Defense Joint Program Committee-6 (JPC-6), Combat Casualty Care Research Program. KRR, RR, and AG were also supported by funds from the Department of Pediatrics and Children’s Research Institute.

### Disclosures

RR is a founder and president of CIAN, Inc, a company that is developing biomarkers for TBI diagnosis and outcome prognostication. Patents and disclosures related to these findings submitted by KRR, RZ, and RR have been licensed to CIAN, Inc.

## Highlights

1. Severe traumatic brain injury acutely reduces cerebral blood flow throughout the brain cortex and results in chronic ischemia that is more focal to the site of impact.
2. Single cell RNA sequencing identified five endothelial cell subclusters, one of which increases in proportion to the others after injury suggesting emergence of an injury response cell phenotype.
3. Traumatic brain injury decreased endothelial cell expression of several cilia genes while prominently upregulating metabolic, hypoxia, and inflammatory molecular pathways.
4. Detection of endothelial cilia protein ARL13B was reduced in brain tissue and increased in the circulation suggesting its potential use as a biomarker of vascular injury.

## Non-standard Abbreviations and Acronyms

CBF: cerebral blood flow
GSEA: gene set enrichment analysis
HBMVECs: human brain microvascular endothelial cells
LSCI: laser speckle contrast imaging
MFI: median fluorescence intensity
scRNAseq: single cell ribonucleic acid sequencing
sTBI: severe traumatic brain injury
UMAP: uniform manifold approximation and projection

Table 1: List of top 10 genes that were most significantly expressed in EC sub-cluster 1.

Table 2: List of ischemia and hypoxia associated genes that were significantly altered in EC cluster post-TBI.

**Supp Fig 1: sTBI causes acute global brain ischemia.** LSCI representative images from one mouse before and up to 120 minutes after sTBI demonstrating the acute changes in cerebral perfusion that occur in both the impacted hemisphere (ipsilateral, bottom) and the non-impacted hemisphere (contralateral, top). Site of impact denoted by red circle.

**Supp Fig 2: Early ischemia transitions to hyperemic CBF in the days following sTBI.** LSCI images from the five mice used to create the data in Figure 1. Brain perfusion images were acquired before sTBI, 30 minutes post sTBI, and 7 days post sTBI. Bright field images of the animals’ perfused brains demonstrate a macroscopic view of the focal contusion injury. Site of impact denoted by red circles.

**Supp Fig 3: sTBI results in chronic focal disturbances in CBF.** LSCI images from two representative mice collected 28 days post sTBI demonstrating the relative hypoperfusion in the impacted hemisphere (bottom) compared to the non-impacted hemisphere (top). Brightfield images demonstrate regions of contusion injuries (black dotted line) that match the LSCI detected ischemia.

**Supp Fig 4: Construction of cDNA libraries from brain-derived single cells for RNA sequencing purpose.** Single cells were cleaned by removing myelin debris, fixed, and processed for library construction as per manufacturer’s (10x Genomics) protocol. For multiplexing purposes, samples were probed by single cell fixed RNA transcriptome probe kit. For a single library construction, 4 samples from each time point were multiplexed by the dual index plate reagent following the preparation of gel emulsion in 10x Genomics chromium X instrument. For the sequencing purpose, 4 libraries were multiplexed. Each library is identifiable by a specific set of i5 and i7 primers as included in the dual index plate. Following the construction as recommended by the manufacturer, cDNA library product was validated in TapeStation (Agilent Technologies), by using High Sensitivity D5000 ScreenTape and analyzing with TapeStation analysis software version 3.1. Specific library products are shown by the red arrows.

**Supp Fig 5: Expression of ischemia markers in EC cluster.** Representative feature plots showing the expression of ischemia markers *S100b, S100a8* and *S100a9* in EC clusters post-TBI.

Supp Fig 6: scRNAseq identifies repertoire of cilia gene expressions in brain derived cells. Top cilia genes that were significantly detectable in any cell cluster (A). Heatmap showing the expression pattern of cilia genes in EC cluster at the indicated time points (B). Top cilia gene expressions that were detectable from EC cluster (C).

## References

1. In: C. Matney, K. Bowman and D. Berwick, eds. *Traumatic Brain Injury: A Roadmap for Accelerating Progress* Washington (DC); 2022.

2. Georges A and J MD. Traumatic Brain Injury *StatPearls* Treasure Island (FL); 2023.

3. Nolan S. Traumatic brain injury: a review. Crit Care Nurs Q. 2005;28:188–94.

4. Salehi A, Zhang JH and Obenaus A. Response of the cerebral vasculature following traumatic brain injury. J Cereb Blood Flow Metab. 2017;37:2320–2339.

5. Martin NA, Patwardhan RV, Alexander MJ, Africk CZ, Lee JH, Shalmon E, Hovda DA and Becker DP. Characterization of cerebral hemodynamic phases following severe head trauma: hypoperfusion, hyperemia, and vasospasm. J Neurosurg. 1997;87:9–19.

6. Honda M, Ichibayashi R, Yokomuro H, Yoshihara K, Masuda H, Haga D, Seiki Y, Kudoh C and Kishi T. Early Cerebral Circulation Disturbance in Patients Suffering from Severe Traumatic Brain Injury (TBI): A Xenon CT and Perfusion CT Study. Neurol Med Chir (Tokyo). 2016;56:501–9.

7. Gaggi NL, Ware JB, Dolui S, Brennan D, Torrellas J, Wang Z, Whyte J, Diaz-Arrastia R and Kim JJ. Temporal dynamics of cerebral blood flow during the first year after moderate-severe traumatic brain injury: A longitudinal perfusion MRI study. Neuroimage Clin. 2023;37:103344.

8. Perez-Barcena J, Romay E, Llompart-Pou JA, Ibanez J, Brell M, Llinas P, Gonzalez E, Merenda A, Ince C and Bullock R. Direct observation during surgery shows preservation of cerebral microcirculation in patients with traumatic brain injury. J Neurol Sci. 2015;353:38–43.

9. Dijkhuizen RM. Advances in MRI-Based Detection of Cerebrovascular Changes after Experimental Traumatic Brain Injury. Transl Stroke Res. 2011;2:524–32.

10. Gardner AJ, Tan CO, Ainslie PN, van Donkelaar P, Stanwell P, Levi CR and Iverson GL. Cerebrovascular reactivity assessed by transcranial Doppler ultrasound in sport-related concussion: a systematic review. Br J Sports Med. 2015;49:1050–5.

11. Len TK and Neary JP. Cerebrovascular pathophysiology following mild traumatic brain injury. Clin Physiol Funct Imaging. 2011;31:85–93.

12. Pop V and Badaut J. A neurovascular perspective for long-term changes after brain trauma. Transl Stroke Res. 2011;2:533-45.

13. Tan CO, Meehan WP, 3rd, Iverson GL and Taylor JA. Cerebrovascular regulation, exercise, and mild traumatic brain injury. Neurology. 2014;83:1665–72.

14. Arneson D, Zhang G, Ahn IS, Ying Z, Diamante G, Cely I, Palafox-Sanchez V, Gomez-Pinilla F and Yang X. Systems spatiotemporal dynamics of traumatic brain injury at single-cell resolution reveals humanin as a therapeutic target. Cell Mol Life Sci. 2022;79:480.

15. Arneson D, Zhang G, Ying Z, Zhuang Y, Byun HR, Ahn IS, Gomez-Pinilla F and Yang X. Single cell molecular alterations reveal target cells and pathways of concussive brain injury. Nat Commun. 2018;9:3894.

16. Bolte AC, Shapiro DA, Dutta AB, Ma WF, Bruch KR, Kovacs MA, Royo Marco A, Ennerfelt HE and Lukens JR. The meningeal transcriptional response to traumatic brain injury and aging. Elife. 2023;12.

17. Garza R, Sharma Y, Atacho DAM, Thiruvalluvan A, Abu Hamdeh S, Jonsson ME, Horvath V, Adami A, Ingelsson M, Jern P, Hammell MG, Englund E, Kirkeby A, Jakobsson J and Marklund N. Single-cell transcriptomics of human traumatic brain injury reveals activation of endogenous retroviruses in oligodendroglia. Cell Rep. 2023;42:113395.

18. Somebang K, Rudolph J, Imhof I, Li L, Niemi EC, Shigenaga J, Tran H, Gill TM, Lo I, Zabel BA, Schmajuk G, Wipke BT, Gyoneva S, Jandreski L, Craft M, Benedetto G, Plowey ED, Charo I, Campbell J, Ye CJ, Panter SS, Nakamura MC, Eckalbar W and Hsieh CL. CCR2 deficiency alters activation of microglia subsets in traumatic brain injury. Cell Rep. 2021;36:109727.

19. Todd BP, Chimenti MS, Luo Z, Ferguson PJ, Bassuk AG and Newell EA. Traumatic brain injury results in unique microglial and astrocyte transcriptomes enriched for type I interferon response. J Neuroinflammation. 2021;18:151.

20. Witcher KG, Bray CE, Chunchai T, Zhao F, O’Neil SM, Gordillo AJ, Campbell WA, McKim DB, Liu X, Dziabis JE, Quan N, Eiferman DS, Fischer AJ, Kokiko-Cochran ON, Askwith C and Godbout JP. Traumatic Brain Injury Causes Chronic Cortical Inflammation and Neuronal Dysfunction Mediated by Microglia. J Neurosci. 2021;41:1597–1616.

21. Zhang L, Yang Q, Yuan R, Li M, Lv M, Zhang L, Xie X, Liang W and Chen X. Single-nucleus transcriptomic mapping of blast-induced traumatic brain injury in mice hippocampus. Sci Data. 2023;10:638.

22. Lee H, Song J, Jung JH and Ko HW. Primary cilia in energy balance signaling and metabolic disorder. BMB Rep. 2015;48:647–54.

23. Nauli SM, Kawanabe Y, Kaminski JJ, Pearce WJ, Ingber DE and Zhou J. Endothelial cilia are fluid shear sensors that regulate calcium signaling and nitric oxide production through polycystin-1. Circulation. 2008;117:1161–71.

24. Steidl ME, Nigro EA, Nielsen AK, Pagliarini R, Cassina L, Lampis M, Podrini C, Chiaravalli M, Mannella V, Distefano G, Yang M, Aslanyan M, Musco G, Roepman R, Frezza C and Boletta A. Primary cilia sense glutamine availability and respond via asparagine synthetase. Nat Metab. 2023;5:385–397.

25. Eisa-Beygi S, Benslimane FM, El-Rass S, Prabhudesai S, Abdelrasoul MKA, Simpson PM, Yalcin HC, Burrows PE and Ramchandran R. Characterization of Endothelial Cilia Distribution During Cerebral-Vascular Development in Zebrafish ( Danio rerio). Arterioscler Thromb Vasc Biol. 2018;38:2806–2818.

26. Diagbouga MR, Morel S, Cayron AF, Haemmerli J, Georges M, Hierck BP, Allemann E, Lemeille S, Bijlenga P and Kwak BR. Primary cilia control endothelial permeability by regulating expression and location of junction proteins. Cardiovasc Res. 2022;118:1583–1596.

27. Dinsmore C and Reiter JF. Endothelial primary cilia inhibit atherosclerosis. EMBO Rep. 2016;17:156–66.

28. Gupta A, Thirugnanam K, Thamilarasan M, Mohieldin AM, Zedan HT, Prabhudesai S, Griffin MR, Spearman AD, Pan A, Palecek SP, Yalcin HC, Nauli SM, Rarick KR, Zennadi R and Ramchandran R. Cilia proteins are biomarkers of altered flow in the vasculature. JCI Insight. 2022;7.

29. Delling M, DeCaen PG, Doerner JF, Febvay S and Clapham DE. Primary cilia are specialized calcium signalling organelles. Nature. 2013;504:311–4.

30. Kochanek PM, Wallisch JS, Bayir H and Clark RSB. Pre-clinical models in pediatric traumatic brain injury-challenges and lessons learned. Childs Nerv Syst. 2017;33:1693–1701.

31. Yu G, Wang LG, Han Y and He QY. clusterProfiler: an R package for comparing biological themes among gene clusters. OMICS. 2012;16:284–7.

32. Vanlandewijck M, He L, Mae MA, Andrae J, Ando K, Del Gaudio F, Nahar K, Lebouvier T, Lavina B, Gouveia L, Sun Y, Raschperger E, Rasanen M, Zarb Y, Mochizuki N, Keller A, Lendahl U and Betsholtz C. A molecular atlas of cell types and zonation in the brain vasculature. Nature. 2018;554:475–480.

33. Kiewitz R, Acklin C, Minder E, Huber PR, Schafer BW and Heizmann CW. S100A1, a new marker for acute myocardial ischemia. Biochem Biophys Res Commun. 2000;274:865–71.

34. McGoron AJ, Capille M, Georgiou MF, Sanchez P, Solano J, Gonzalez-Brito M and Kuluz JW. Post traumatic brain perfusion SPECT analysis using reconstructed ROI maps of radioactive microsphere derived cerebral blood flow and statistical parametric mapping. BMC Med Imaging. 2008;8:4.

35. Pasco A, Lemaire L, Franconi F, Lefur Y, Noury F, Saint-Andre JP, Benoit JP, Cozzone PJ and Le Jeune JJ. Perfusional deficit and the dynamics of cerebral edemas in experimental traumatic brain injury using perfusion and diffusion-weighted magnetic resonance imaging. J Neurotrauma. 2007;24:1321–30.

36. Kramer DR, Winer JL, Pease BA, Amar AP and Mack WJ. Cerebral vasospasm in traumatic brain injury. Neurol Res Int. 2013;2013:415813.

37. Martin RM, Wright MJ, Lutkenhoff ES, Ellingson BM, Van Horn JD, Tubi M, Alger JR, McArthur DL and Vespa PM. Traumatic hemorrhagic brain injury: impact of location and resorption on cognitive outcome. J Neurosurg. 2017;126:796–804.

38. Xia F, Keep RF, Ye F, Holste KG, Wan S, Xi G and Hua Y. The Fate of Erythrocytes after Cerebral Hemorrhage. Transl Stroke Res. 2022;13:655–664.

39. Xu L, Nirwane A, Xu T, Kang M, Devasani K and Yao Y. Fibroblasts repair blood-brain barrier damage and hemorrhagic brain injury via TIMP2. Cell Rep. 2022;41:111709.

40. Bouma GJ, Muizelaar JP, Choi SC, Newlon PG and Young HF. Cerebral circulation and metabolism after severe traumatic brain injury: the elusive role of ischemia. J Neurosurg. 1991;75:685–93.

41. Hiler M, Czosnyka M, Hutchinson P, Balestreri M, Smielewski P, Matta B and Pickard JD. Predictive value of initial computerized tomography scan, intracranial pressure, and state of autoregulation in patients with traumatic brain injury. J Neurosurg. 2006;104:731–7.

42. Lang EW, Lagopoulos J, Griffith J, Yip K, Mudaliar Y, Mehdorn HM and Dorsch NW. Noninvasive cerebrovascular autoregulation assessment in traumatic brain injury: validation and utility. J Neurotrauma. 2003;20:69–75.

43. Rohde D, Schon C, Boerries M, Didrihsone I, Ritterhoff J, Kubatzky KF, Volkers M, Herzog N, Mahler M, Tsoporis JN, Parker TG, Linke B, Giannitsis E, Gao E, Peppel K, Katus HA and Most P. S100A1 is released from ischemic cardiomyocytes and signals myocardial damage via Toll-like receptor 4. EMBO Mol Med. 2014;6:778–94.

44. Tanaka Y, Shigemura N, Kawamura T, Noda K, Isse K, Stolz DB, Billiar TR, Toyoda Y, Bermudez CA, Lyons-Weiler J and Nakao A. Profiling molecular changes induced by hydrogen treatment of lung allografts prior to procurement. Biochem Biophys Res Commun. 2012;425:873–9.

45. Wang W, McKinnie SM, Patel VB, Haddad G, Wang Z, Zhabyeyev P, Das SK, Basu R, McLean B, Kandalam V, Penninger JM, Kassiri Z, Vederas JC, Murray AG and Oudit GY. Loss of Apelin exacerbates myocardial infarction adverse remodeling and ischemia-reperfusion injury: therapeutic potential of synthetic Apelin analogues. J Am Heart Assoc. 2013;2:e000249.

46. Wu Y, Wang X, Zhou X, Cheng B, Li G and Bai B. Temporal Expression of Apelin/Apelin Receptor in Ischemic Stroke and its Therapeutic Potential. Front Mol Neurosci. 2017;10:1.

47. Krock BL, Skuli N and Simon MC. Hypoxia-induced angiogenesis: good and evil. Genes Cancer. 2011;2:1117–33.

48. Lin X, Chen L, Jullienne A, Zhang H, Salehi A, Hamer M, T CH, Obenaus A and Xu X. Longitudinal dynamics of microvascular recovery after acquired cortical injury. Acta Neuropathol Commun. 2022;10:59.

49. Park E, Bell JD, Siddiq IP and Baker AJ. An analysis of regional microvascular loss and recovery following two grades of fluid percussion trauma: a role for hypoxia-inducible factors in traumatic brain injury. J Cereb Blood Flow Metab. 2009;29:575–84.

50. Hay JR, Johnson VE, Young AM, Smith DH and Stewart W. Blood-Brain Barrier Disruption Is an Early Event That May Persist for Many Years After Traumatic Brain Injury in Humans. J Neuropathol Exp Neurol. 2015;74:1147–57.

51. Tang J, Kang Y, Huang L, Wu L and Peng Y. TIMP1 preserves the blood-brain barrier through interacting with CD63/integrin beta 1 complex and regulating downstream FAK/RhoA signaling. Acta Pharm Sin B. 2020;10:987–1003.

52. Tang J, Kang Y, Zhou Y, Shang N, Li X, Wang H, Lan J, Wang S, Wu L and Peng Y. TIMP2 ameliorates blood-brain barrier disruption in traumatic brain injury by inhibiting Src-dependent VE-cadherin internalization. J Clin Invest. 2023;134.

53. Thirugnanam K, Prabhudesai S, Van Why E, Pan A, Gupta A, Foreman K, Zennadi R, Rarick KR, Nauli SM, Palecek SP and Ramchandran R. Ciliogenesis mechanisms mediated by PAK2-ARL13B signaling in brain endothelial cells is responsible for vascular stability. Biochem Pharmacol. 2022;202:115143.

54. Schneider ALC, Huie JR, Jain S, Sun X, Ferguson AR, Lynch C, Yue JK, Manley GT, Wang KKW, Sandsmark DK, Campbell C and Diaz-Arrastia R. Associations of Microvascular Injury-Related Biomarkers With Traumatic Brain Injury Severity and Outcomes: A Transforming Research and Clinical Knowledge in Traumatic Brain Injury (TRACK-TBI) Pilot Study. J Neurotrauma. 2023;40:1625–1637.

55. Bazarian JJ, Biberthaler P, Welch RD, Lewis LM, Barzo P, Bogner-Flatz V, Gunnar Brolinson P, Buki A, Chen JY, Christenson RH, Hack D, Huff JS, Johar S, Jordan JD, Leidel BA, Lindner T, Ludington E, Okonkwo DO, Ornato J, Peacock WF, Schmidt K, Tyndall JA, Vossough A and Jagoda AS. Serum GFAP and UCH-L1 for prediction of absence of intracranial injuries on head CT (ALERT-TBI): a multicentre observational study. Lancet Neurol. 2018;17:782–789.

