## Supplementary figures and images for "Severe traumatic brain injury temporally affects cerebral blood flow, endothelial cell phenotype, and cilia"

### Supplemental figures

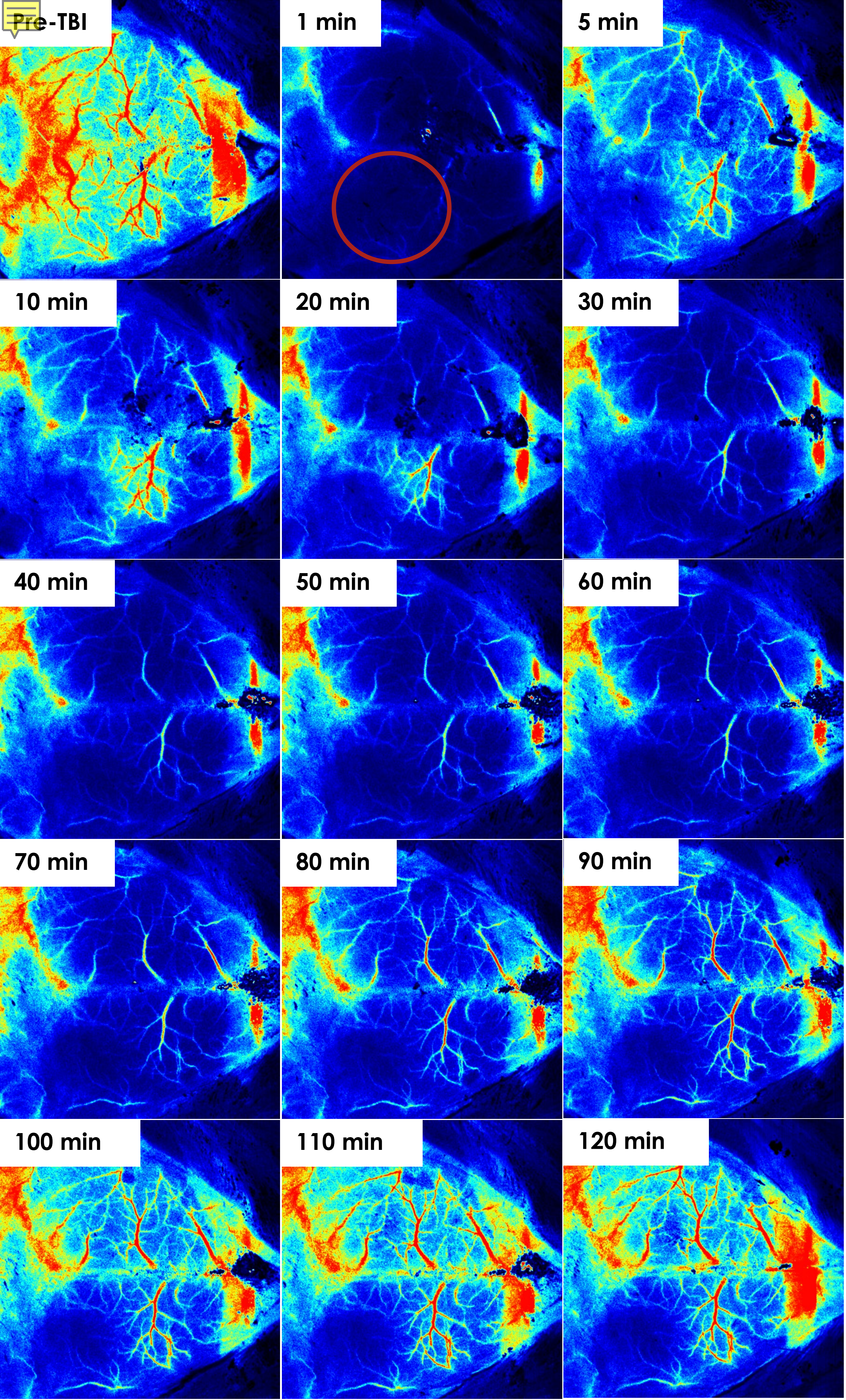

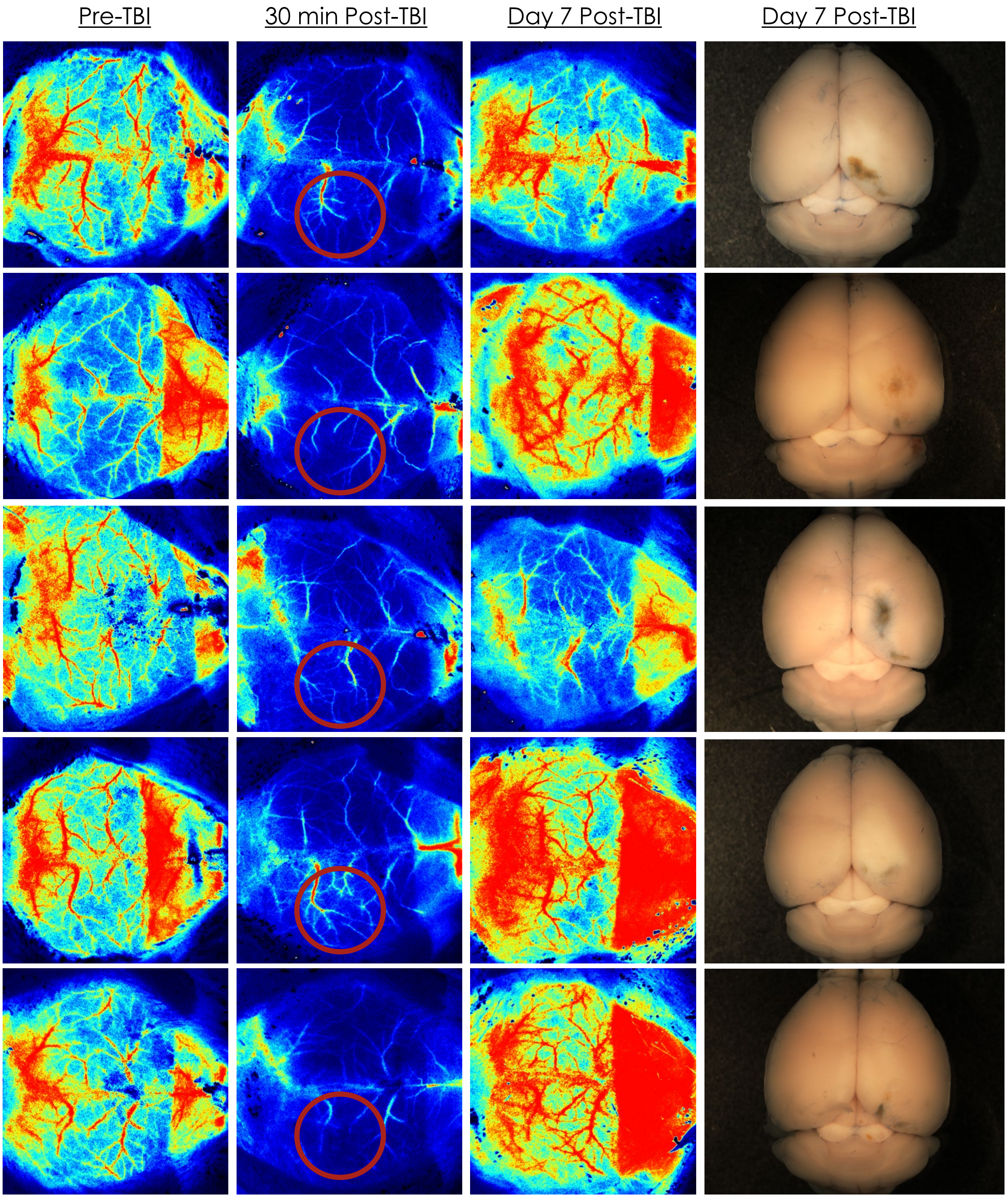

Day 28 Post sTBI

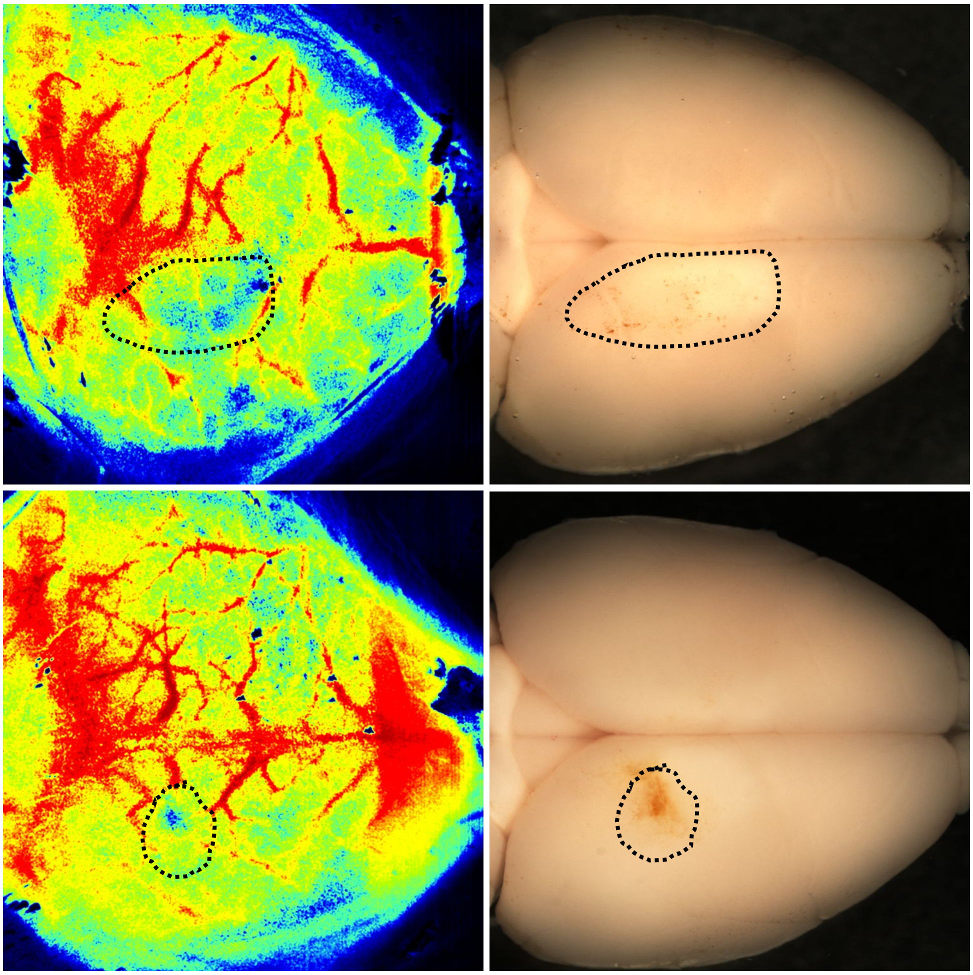

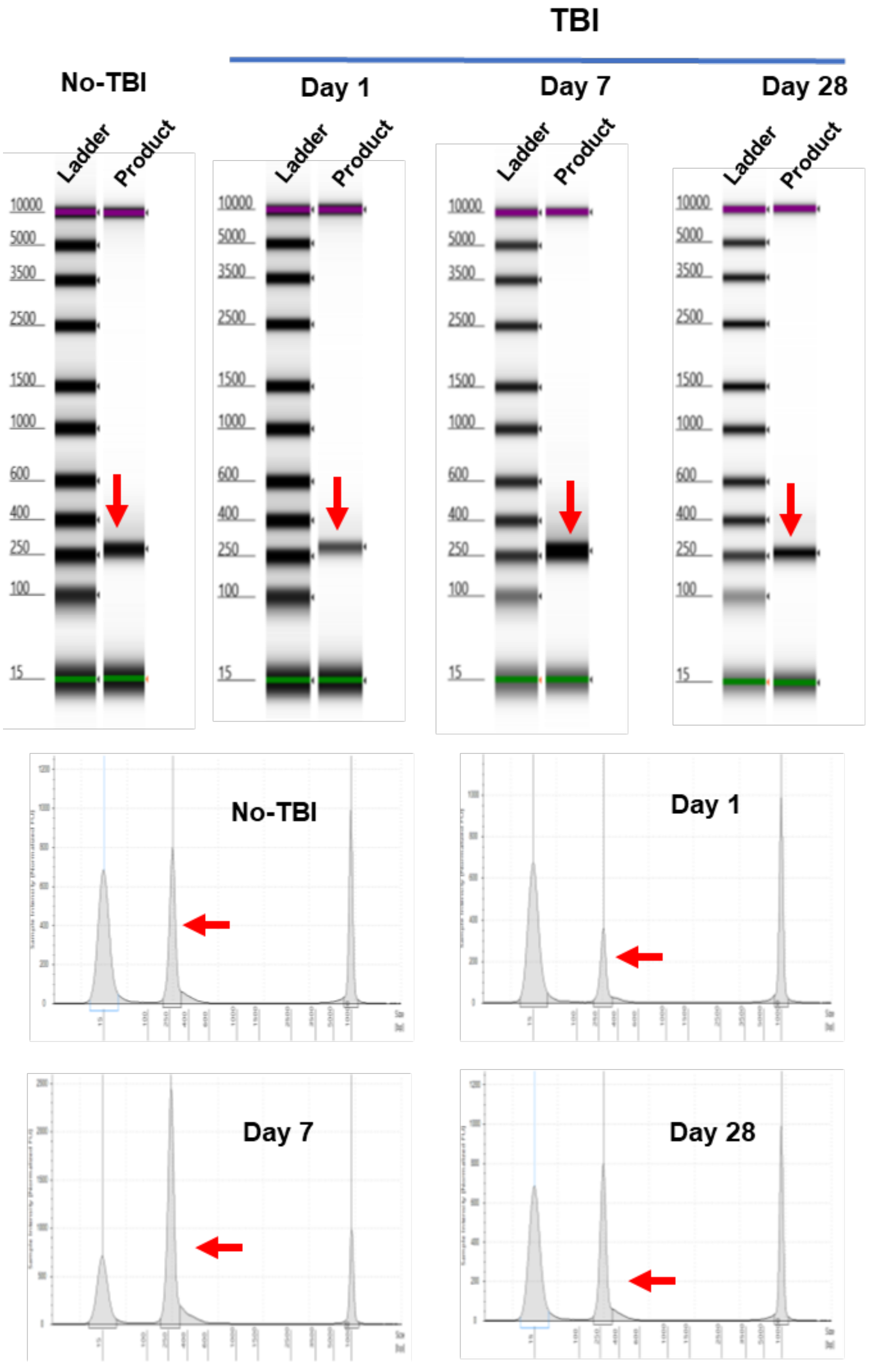

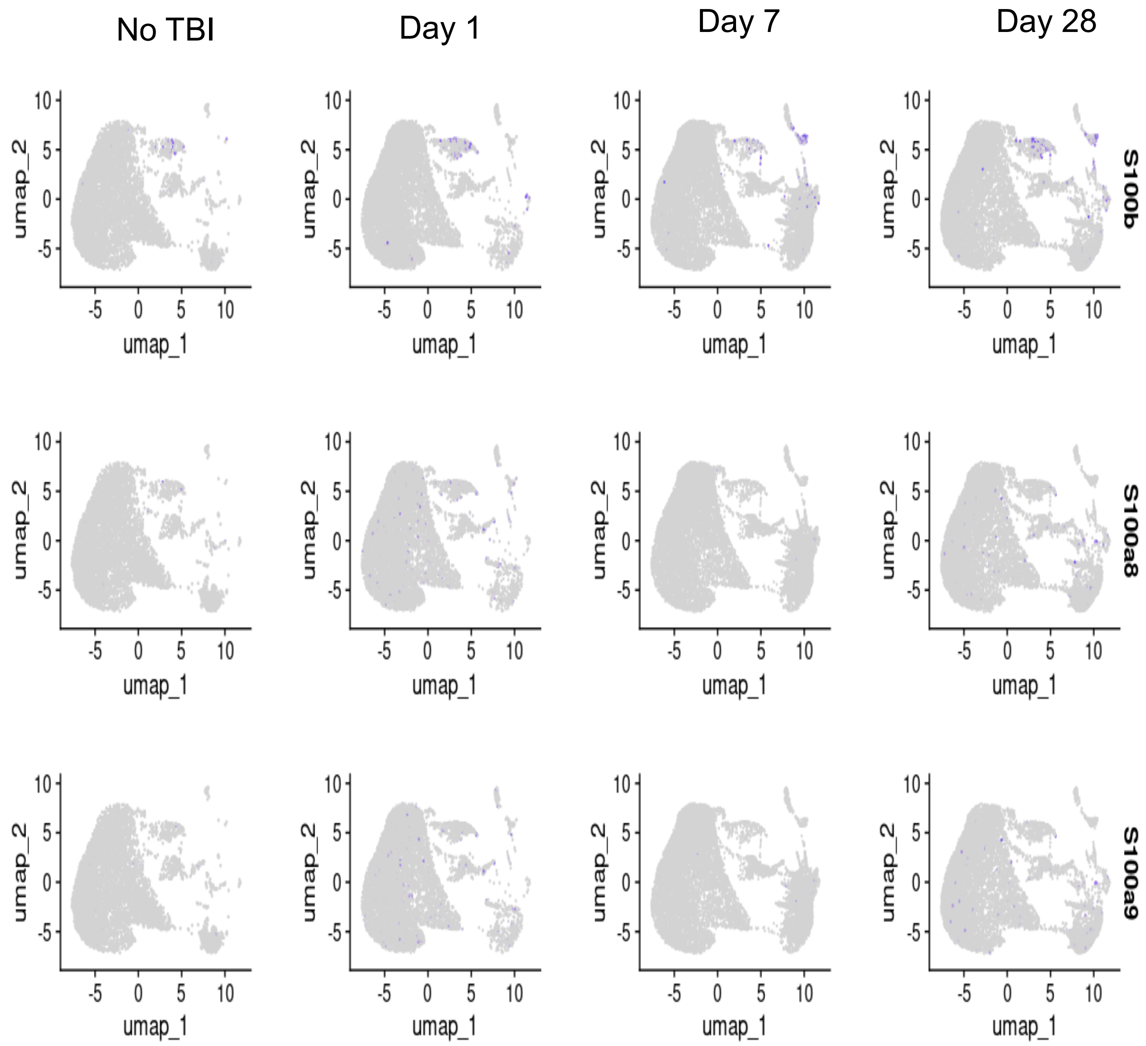

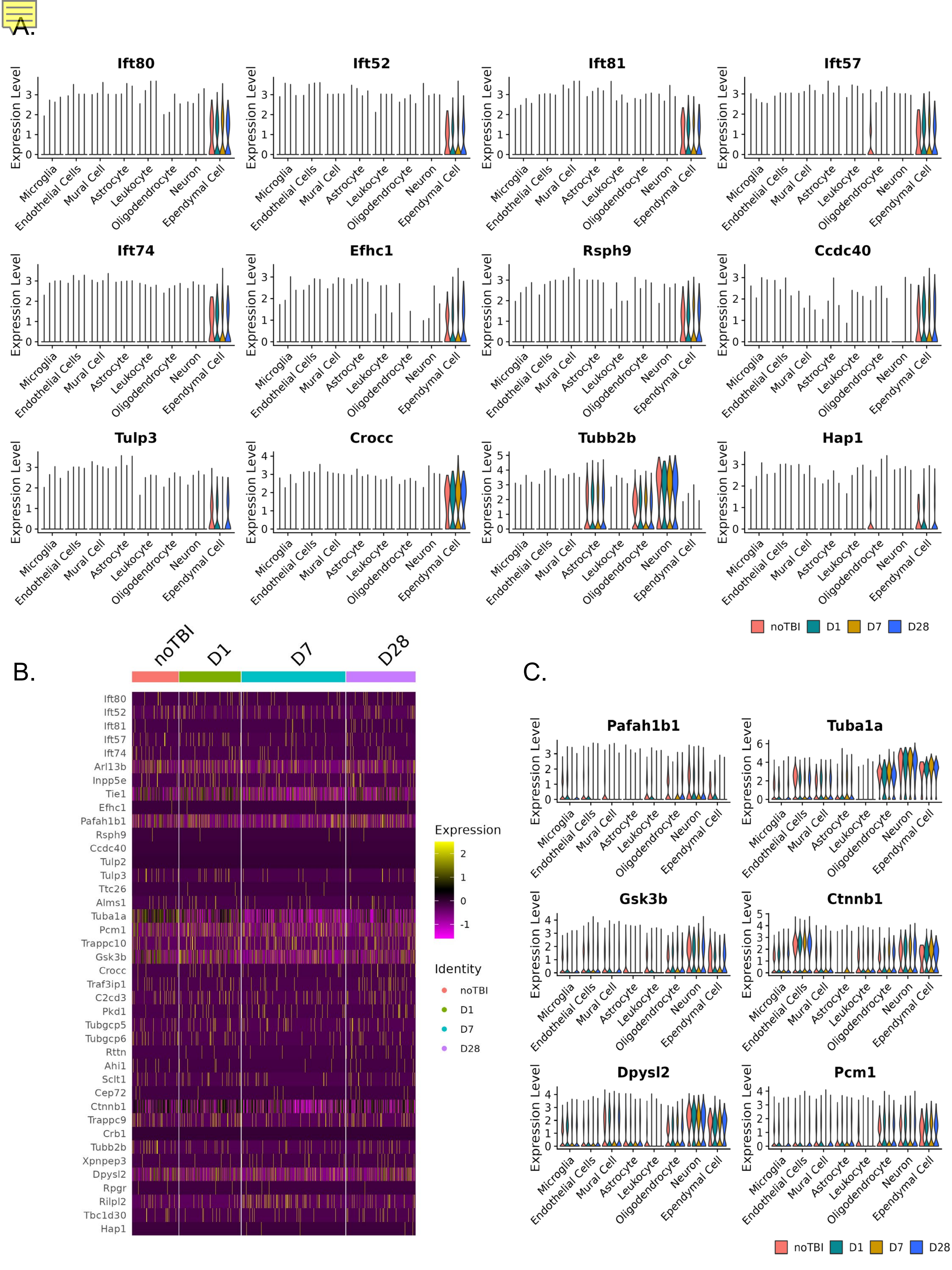
